# Kinetically distinct phases of tau on microtubules regulate kinesin motors and severing enzymes

**DOI:** 10.1101/424374

**Authors:** Valerie Siahaan, Jochen Krattenmacher, Amayra Hernandez-Vega, Anthony A. Hyman, Stefan Diez, Zdenek Lansky, Marcus Braun

## Abstract

Tau is an intrinsically disordered protein, which diffuses on microtubules. In neurodegenerative diseases collectively termed tauopathies, tau malfunction and its detachment from axonal microtubules is correlated with microtubule degradation. It is known that tau can protect microtubules from microtubule-degrading enzymes, such as katanin. However, how tau can fulfill such regulative function is still unclear. Using in vitro reconstitution, we here show that tau molecules on microtubules cooperatively form islands of an ordered layer with regulatory qualities distinct from a comparably dense layer of diffusible tau. These islands shield the microtubules from katanin and kinesin-1 but are penetrable by kinesin-8 which causes the islands to disassemble. Our results indicate a new phase of tau, constituting an adjustable protective sheath around microtubules.

## Introduction

Tau is an intrinsically disordered microtubule associated protein (MAP), which localizes preferentially to neuronal axons and is involved in a number of neurodegenerative diseases (*1*-*3*). Tau diffuses along microtubules (*4*) and can enhance the stability of microtubules both, by its presence (*5*) and by its capability to regulate the interaction of other MAPs with the microtubule surface. For example, displacement of tau from the axon is one of the hallmark events during the onset of the Alzheimer’s disease (*6*). Tau mislocalization leaves axonal microtubules unprotected against microtubule-severing enzymes, such as katanin (*7*), which leads to their destabilization and the eventual degeneration of the axon. Tau can also regulate the microtubule interactions of other MAPs, such as molecular motors (*8*-*13*). However, the underlying molecular mechanism of how tau as an intrinsically disordered protein can fulfill such regulative functions is still unknown. Here we study how the collective behaviour of tau can influence the activity of motors and severing proteins on microtubules. Using in vitro reconstitution assays, we found that on the microtubule surface tau can form islands of cohesive deposits in a pool of diffusible tau molecules. We observed that the islands prevented the interaction of katanin and kinesin-1 motors with the microtubule surface. In contrast, super-processive kinesin-8 motors, initially moving in the low-density regions of diffusible tau molecules, could penetrate the islands where they caused island disassembly. Our results show that two phases of tau, co-existing on the microtubule surface, differentially regulate the interactions of MAPs with the microtubule surface.

## Results

### Tau on microtubules separates into two kinetically distinct phases

To study the interaction of tau with microtubules, we immobilized Atto-647-labeled microtubules on a coverslip, added (full-length) tau labeled with monomeric-enhanced green fluorescent protein (tau-mEGFP) and performed time-lapse imaging using TIRF microscopy (Fig. 1A, Methods). After the addition of 20 nM tau-mEGFP we observed the formation of high-density tau-mEGFP regions on the microtubules (further on termed ‘islands’) surrounded by regions of low-density tau-mEGFP (Fig. 1B,C,D, Movie S1). Tau-mEGFP islands (i) were nucleated from diffraction-limited spots, (ii) appeared (70% of them) within 20 s after the addition of tau (Fig. S1A), (iii) grew non-uniformly at their boundaries to highly variable lengths (Fig. S1B), and (iv) covered 17 ± 8 % of the total available microtubule length after 15 min. Quantification of the tau-mEGFP density on the microtubules yielded 0.09 ± 0.04 nm^-1^ in the low-density regions and 0.48 ± 0.09 nm^-1^ in the islands (Methods). The tau-mEGFP density in the islands stayed constant during the growth phase (Fig. 1E), suggesting that the islands grow predominantly by the addition of tau molecules at their boundaries. When the boundaries of growing islands collided, they merged with neighboring growing islands (Fig. 1C). At these instances, we never observed an increase in the tau-mEGFP density, suggesting that each island itself already occupies the entire accessible surface of the microtubule. After removal of tau-mEGFP from solution (Fig. 1F, Movie S2) tau-mEGFP outside the islands unbound from the microtubules uniformly along the whole length of the low-density regions within a few seconds (Fig. S1C). Islands, by contrast, persisted over several minutes. Strikingly, in the absence of tau-mEGFP in solution, the islands disassembled from their boundaries until they fully disappeared, with the island boundaries receding with an average velocity of 6.6 ± 5.2 nm/s. During disassembly, the islands occasionally underwent fission (Fig. 1F). We determined the mean residence time of tau-mEGFP inside the disassembling islands to be 93 ± 6 min (Fig. 1G, S1C), showing that the islands disassembled predominantly at their boundaries. This cohesive quality of the tau islands, together with their (dis)assembly kinetics, suggest that tau molecules cooperate in order to form the islands. Because the N-terminus of tau has been reported to be important for tau-tau interactions (*14*), we repeated the described experiments with a truncated tau construct comprising the four-repeat microtubule-binding domain and the C-terminus but lacking the N-terminus (tau∆N-mEGFP, Fig. S1D). Although tau∆N-mEGFP did interact with the microtubules, we did not observe any island formation even at tau∆N-mEGFP concentrations as high as 0.5 µM. To test the robustness of the island assembly, we varied the ionic strength of the assay buffer and observed the islands to assemble over a broad range of conditions (0 - 125 mM KCl in the assay buffer, Methods). Combined, our results show that full-length tau on microtubules can separate into two kinetically distinct phases.

**Fig. 1.**
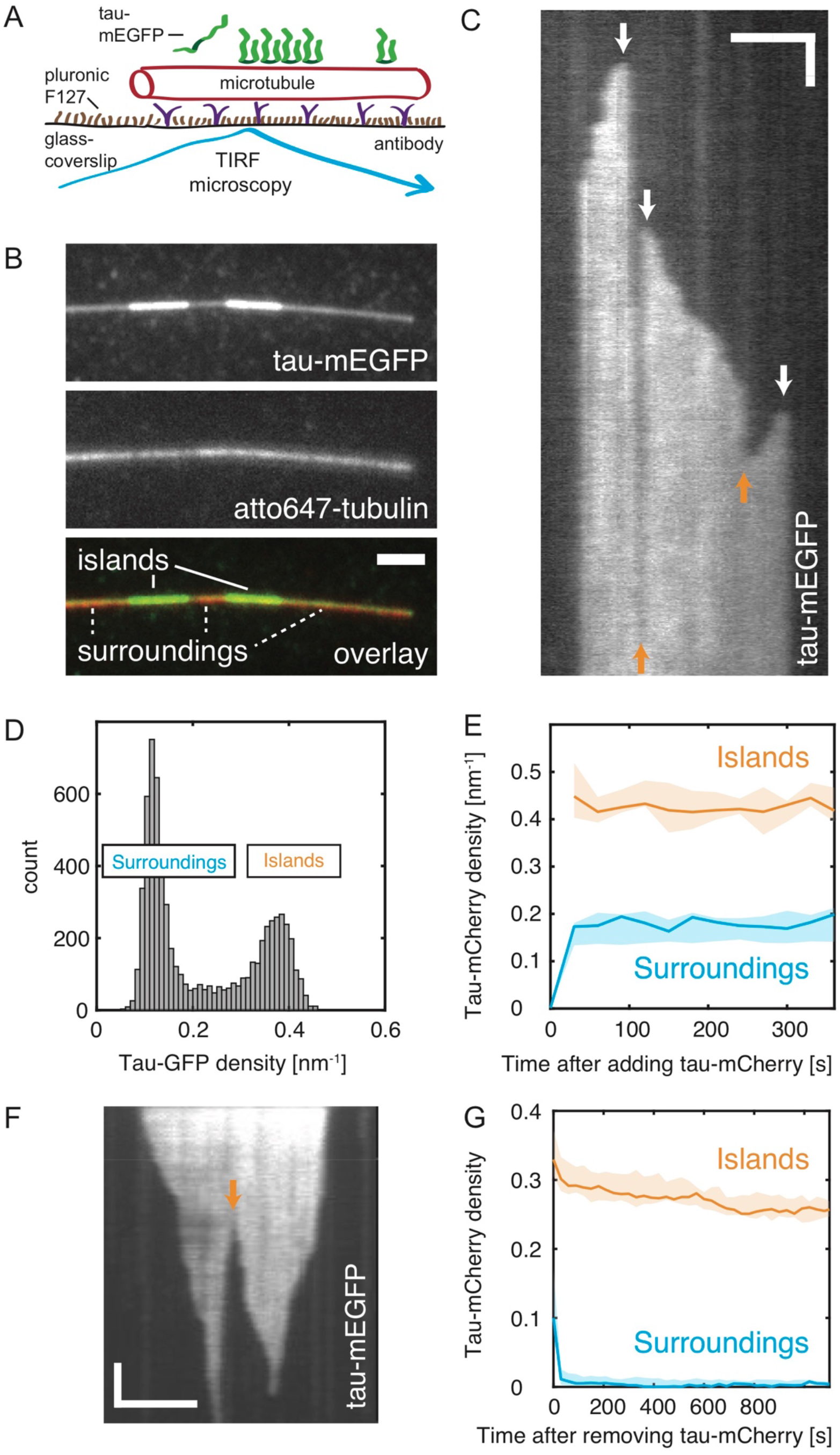
Tau forms dynamic high-density islands on microtubules. **(A)** Schematics depicting the assay geometry. **(B)** Multichannel fluorescence micrograph showing the islands of high tau-mEGFP density (bright green) surrounded by regions of low tau-mEGFP density (dim green) on the Atto-647-labeled microtubule (red). **(C)** Kymograph showing the fluorescence signal of tau-mEGFP on the microtubule after the addition 20 nM tau-mEGFP. Initially the microtubule is covered by low tau-mEGFP density, over time high density tau-mEGFP islands get nucleated and grow. White arrows indicate the nucleation points. Orange arrows indicate merging of two neighboring islands growing towards each other. **(D)** Distribution of fluorescence intensity of tau-mEGFP along the microtubules such as shown in (C) showing two distinct populations. **(E)** The tau-mEGFP density within and outside of the islands after the addition of tau-mEGFP. The tau-mEGFP density in the island regions is constant during the formation of the islands. This experiment was performed with lower frame rate than in (C) to minimize photo-bleaching (Methods). **(F)** Kymograph showing the fluorescence signal of tau-mEGFP on the microtubule during the disassembly of the islands after the removal of tau-mEGFP from solution. Orange arrow indicates fission events during island disassembly. **(G)** The tau-mEGFP density within and outside the islands after the removal of tau-mEGFP from solution. Outside the islands, the drop in the density corresponds to the mean residence time of tau-mEGFP on the microtubule of 1.9 ± 0.2 s, within the islands the mean residence time was 93 ± 20 min (for fits see Fig. S1C). This experiment was performed with lower frame rate than (F) to minimize photo-bleaching (Methods). All scale bars, vertical 5 s, horizontal 2 µm.

### Tau molecules in the islands are stationary but exchange with tau in solution

To further explore the dynamics of tau molecules in the islands we formed islands using 20 nM mCherry-labelled tau (tau-mCherry) and, after 15 minutes, replaced the assay buffer by a solution containing 20 nM tau-mEGFP (Fig. 2A, Movie S3). In the low-density regions tau-mCherry rapidly dissociated from the microtubules with an average time constant of 2.2 s. This value is comparable to the 1.9 s estimated for the situation discussed in Fig. 1, where tau-mEGFP was completely removed from solution. By contrast, within the islands tau-mCherry dissociated markedly slower (average time constant of 31 s, Fig. S2A, S2B) but substantially faster than in the situation when tau was completely removed from solution (compare to Fig. 1G, S1C). The latter observation suggests that tau unbinding from the islands depends on the tau concentration in solution (Fig. 2B) highlighting the multivalency of the interaction in analogy to the concentration-dependent unbinding rates of DNA-binding proteins (*15*). Simultaneously, in exchange for the leaving tau-mCherry, tau-mEGFP molecules bound rapidly to the low-density regions and with a slower time constant into the islands (Fig. 2A, S2A). Turnover from tau-mCherry to tau-mEGFP occurred uniformly along the whole lengths of both regions. In addition, and similar to the initial formation of tau-mCherry islands, tau-mEGFP also associated with the boundaries of existing islands, elongating them (Fig. 2A). To further study the spatio-temporal dynamics of tau in the high-density and low-density regions, we formed islands using 20 nM tau-mCherry and 1 nM tau-mEGFP. This strategy allowed us to observe the motion of individual tau-mEGFP molecules in a surrounding dominated by tau-mCherry molecules. In the low-density regions, single tau-mEGFP molecules diffused rapidly (Fig. 2C). By contrast, in the islands the tau-mEGFP molecules were bound stationarily to the microtubule surface (Fig. 2C). Occasionally, single tau-mEGFP molecules initially diffusing outside an island became stationary when associating with an island boundary (Fig. 2C). Combined, our results show that the tau molecules localizing in the islands are stationary, but can exchange with tau in solution.

**Fig. 2.**
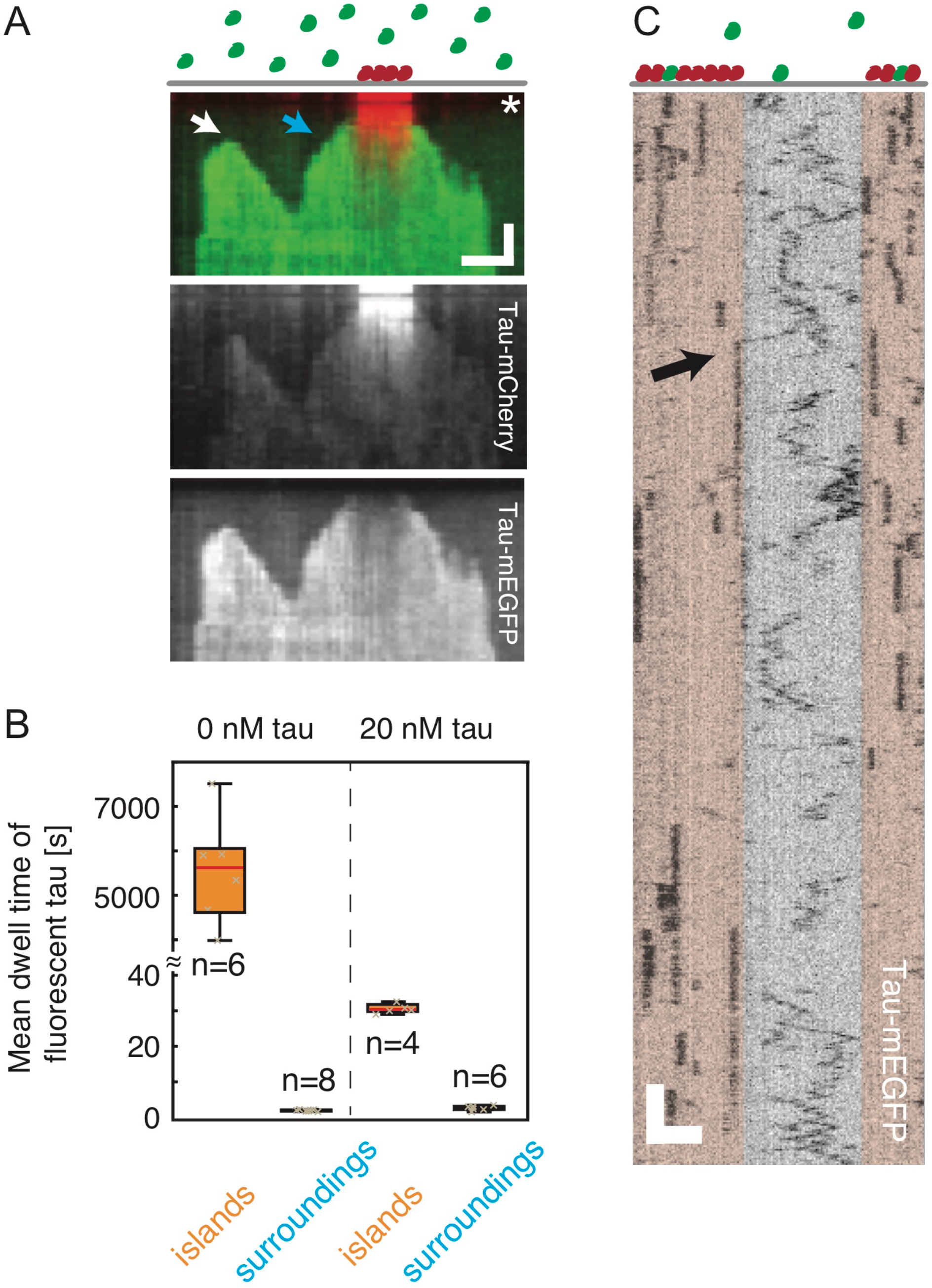
Tau molecules in the islands are stationary but exchange with tau in solution. **(A)** Multichannel kymograph showing the islands pre-formed in presence of 20 nM tau-mCherry (red). After the addition (time marked by white asterisk) of 20 nM tau-mEGFP (green), removing most of the tau-mCherry from solution we observed the exchange of tau-mCherry for tau-mEGFP within and outside of the islands. This exchange occurred along the whole length of the island regions. Additionally, islands resumed their growth by the addition of tau-mEGFP to their boundaries (marked by blue arrow). New islands were also nucleated (marked by white arrow). Scale bars, vertical 20 s, horizontal 2 µm. **(B)** Box-plot of the dwell times of fluorescently labeled tau outside and inside the islands in the absence of tau in solution (data from Fig. 1F,G) and in the presence of 20 nM tau-mEGFP in solution (data from (A)). **(C)** Intensity-inverted kymograph showing single tau-mEGFP molecules interacting with a microtubule crowded with tau-mCherry. The position of the islands are indicated by beige transparent boxes overlayed with the kymograph. Outside of the islands, tau-mEGFP diffuses rapidly, whereas inside the islands, tau-mEGFP is bound stationarily. Occasionally, we observed a diffusing tau-mEGFP molecule stopping while getting associated with an island (black arrow). Scale bars, vertical 1 s, horizontal 2 µm.

### Tau islands regulate the interaction of MAPs with microtubules

To investigate whether the formation of tau islands affects the interaction of other proteins with microtubules, we formed tau islands using tau-mCherry and tested the interaction of the following MAPs (in the presence of 1 mM ATP) with such tau-mCherry-decorated microtubules: i) Processive microtubule-transport motor kinesin-1: After addition of kinesin-1 (truncated rat kinesin rKin-430-GFP, concentration of 60 nM) to microtubules in the presence of 20 nM tau-mCherry, we observed single motors moving processively through the low-density tau regions outside the islands. However, when reaching the boundary of a tau island, the motors dissociated instantaneously from the microtubule (Fig. 3A). No motors bound to the tau islands from solution, resulting in rKin-430-GFP localizing solely to the regions outside of the islands. These results show that tau islands prevent kinesin-1-mediated transport along the microtubule surface. ii) Microtubule-severing enzyme katanin: After the addition of katanin (mouse katanin p60/p80C-strep-GFP (*16*), concentration of 0.1 µM) to microtubules in the presence of 20 nM tau-mCherry, we observed katanin-GFP-mediated microtubule severing with subsequent microtubule disassembly exclusively in the low-density tau regions outside the islands (Fig. 3B, Movie S4). Only on longer time scales, the island-decorated parts of the microtubules started to disassemble predominantly from their boundaries (Fig. 3B, Movie S4). These results show that tau islands can form a protective sheath on the microtubule surface hindering the activity of enzymes altering the microtubule structure, such as microtubule-severing factors. iii) Super-processive microtubule-depolymerising motor kinesin-8: After the addition of kinesin-8 (full-length yeast kip3-GFP, concentration of 45 nM) to microtubules in the presence of 10 nM tau-mCherry, we observed that kip3-GFP molecules could move both, in the low-density tau regions and, at a decreased velocity, in the islands (Fig. 3C,D, S3A Movie S5). Interestingly, kip3-GFP (which was previously shown to form traffic jams at the ends of highly-stabilized microtubules (*17*)) accumulated at the boundaries of the tau islands and caused the enhanced unbinding of tau-mCherry at these positions, eventually leading to the complete removal of the islands (Fig. 3C,D, Movie S5). These results show that not only do tau islands regulate the interaction of other MAPs with the microtubule surface but that, vice versa the activity of other MAPs, such as highly processive motors proteins, can also regulate the dynamics of the islands themselves.

**Fig. 3.**
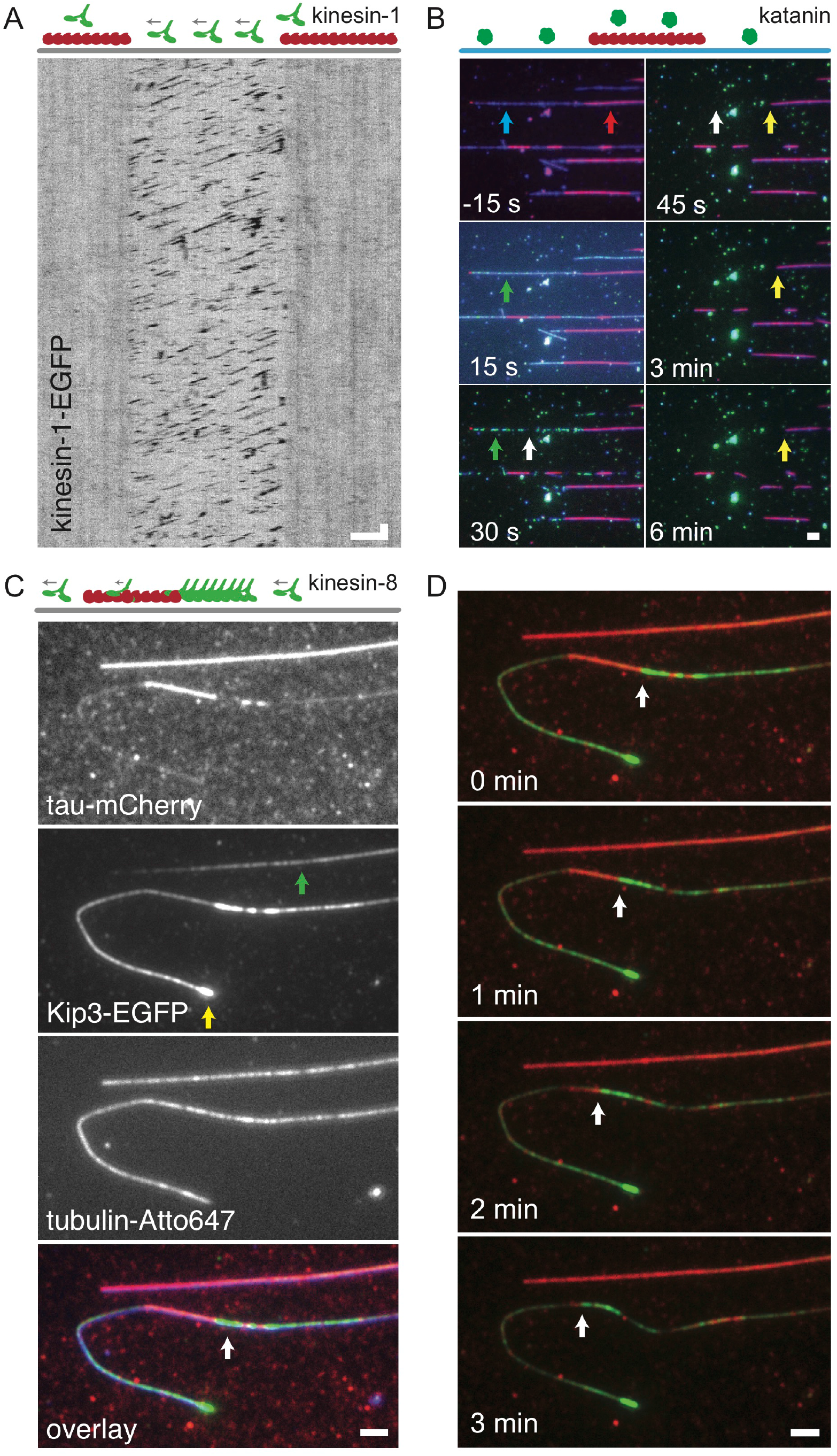
Co-existing tau phases differentially regulate the interaction of MAPs with microtubules. **(A)** Intensity-inverted kymograph showing kinesin-1-GFP molecules bind to and move processively outside of the islands (the positions of the islands are indicated by schematics above the kymograph). We observed no kinesin-1-GFP binding within the island regions. When reaching the island boundaries, the kinesin-1-GFP molecules immediately dissociate from the microtubule. **(B)** Fluorescence micrographs showing the katanin-GFP-driven (green, position indicated by green arrow) severing of Atto-647-microtubules (blue) decorated with tau-mCherry islands (red, position indicated by red arrow) interspersed by regions of low tau-mEGFP density (indicated by blue arrow). Initially, microtubule severing and microtubule disassembly occurred only in the regions outside of the islands (white arrows). On longer time scales (after approximately 1 minute), when the microtubule regions not protected by the islands already disassembled, katanin-GFP induced shortening of the island-covered regions of the microtubule (yellow arrows). **(C)** Multichannel fluorescence micrographs showing that kip3-GFP (kinesin-8, green) localizes in both regions, outside and within the tau-mCherry (red) islands (indicated by green arrow) on Atto-647-labeled microtubules (blue) and accumulates at the microtubule ends (yellow arrow) and in front of the islands (white arrow). **(D)** Multicolor timelapse micrographs showing that kip3-GFP (green) accumulating in front of the tau-mCherry island (red), can remove the island by displacing the tau-mCherry from the island edge (receding of the island boundary indicated by white arrow). All scale bars, vertical 2 s, horizontal 2 µm.

### A rapidly turning-over tau phase can colocalize with the islands

Thus far, in our experiments we used low nM concentrations of tau, sufficient to form the described islands. However, physiological tau concentrations are in the range of 0.5 - 2 µM (*18*). We therefore asked, how island formation and dynamics would be affected by such elevated tau concentrations. When adding 0.8 µM tau-mCherry to surface-immobilized microtubules, we found that the microtubules were uniformly covered with tau at a density of 0.7 ± 0.1 nm^-1^, significantly higher than the island density presented in Fig. 1. Upon removal of tau-mCherry from solution, most of the tau unbound from the microtubules within a few seconds. Only then, the tau islands became visible and disassembled in a similar manner as presented in Fig. 1F and G (Fig. 4A,B). Strikingly, already 30 s after tau-mCherry removal the tau density inside the disassembling islands had lowered to 0.26 ± 0.04 nm^-1^, comparable to the density in the islands described in Fig. 1. To systematically explore how the formation of tau islands depends on the tau concentration in solution we performed an experiment in which we repeated cycles of i) forming tau islands at increasing concentrations of tau-mEGFP in solution followed by ii) completely removing tau-mEGFP from solution. Below tau-mEGFP concentrations of 5 nM we did not observe any island formation. Above this critical concentration, the fraction of the microtubule length covered by the islands increased with increasing tau concentration (Fig. S4A). Likewise, the tau-mEGFP density outside the islands increased with increasing tau-mEGFP concentration. After each tau-mEGFP removal, the density in these regions returned to the background level within several seconds (Fig. 4C, S4B). Interestingly, the tau-mEGFP density in the islands also increased with increasing tau-mEGFP concentration in solution. However, in contrast to the regions outside of the islands, after removal of tau-mEGFP, the density in the islands decayed in two phases (Fig. 4D, S4B). Within a few seconds after tau-mEGFP removal, a fast density drop occurred uniformly along the whole length of the islands followed by a slow density decrease as observed in Fig. 1G. In all experiments, density in the islands after the fast drop was in the range between 0.25 and 0.45 nm^-1^, independent of the initial tau concentration in solution (Fig. 4D, S4B). This suggests that at elevated tau concentrations, two different populations of tau can co-localize within the island regions: i) tau molecules, which bind to the microtubule at the above mentioned characteristic density (0.25 - 0.45 nm^-1^) independent of the tau concentration in solution and which exhibit slow characteristic turn-over presented in Figs. 1 and 2 and ii) tau molecules, which turn over rapidly (likely not involved in cooperative island formation) and whose density depends on the tau concentration in solution, similar to the density in the regions outside the islands (Fig. S4C). Forming the islands using tau-mCherry concentrations as high as 1.5 µM showed that the tau-mCherry density on the microtubule reached saturation (Fig. S4D) as reported before (*19*), indicating that in both regions tau associates with a finite number of interaction sites on the microtubule surface and that all tau molecules are in direct contact with the microtubule lattice. Finally, we wondered whether high-density (~ 0.4 nm^-1^ and higher) tau in the phase that rapidly turns over (i.e. outside the islands) can constitute a protective layer on the microtubules similarly to the islands. We thus formed islands using 0.8 µM tau-mCherry and exposed them to 0.2 µM katanin-GFP similarly to the experiment presented in Fig. 3B. Strikingly, we observed qualitatively identical results, namely protection in the islands but severing and microtubule disassembly outside the islands (Fig. 4E, Movie 6). This experiment shows that it is not the density of tau on the microtubule surface, but the presence of the tau phase that slowly turns over, which protects the microtubule.

**Fig. 4.**
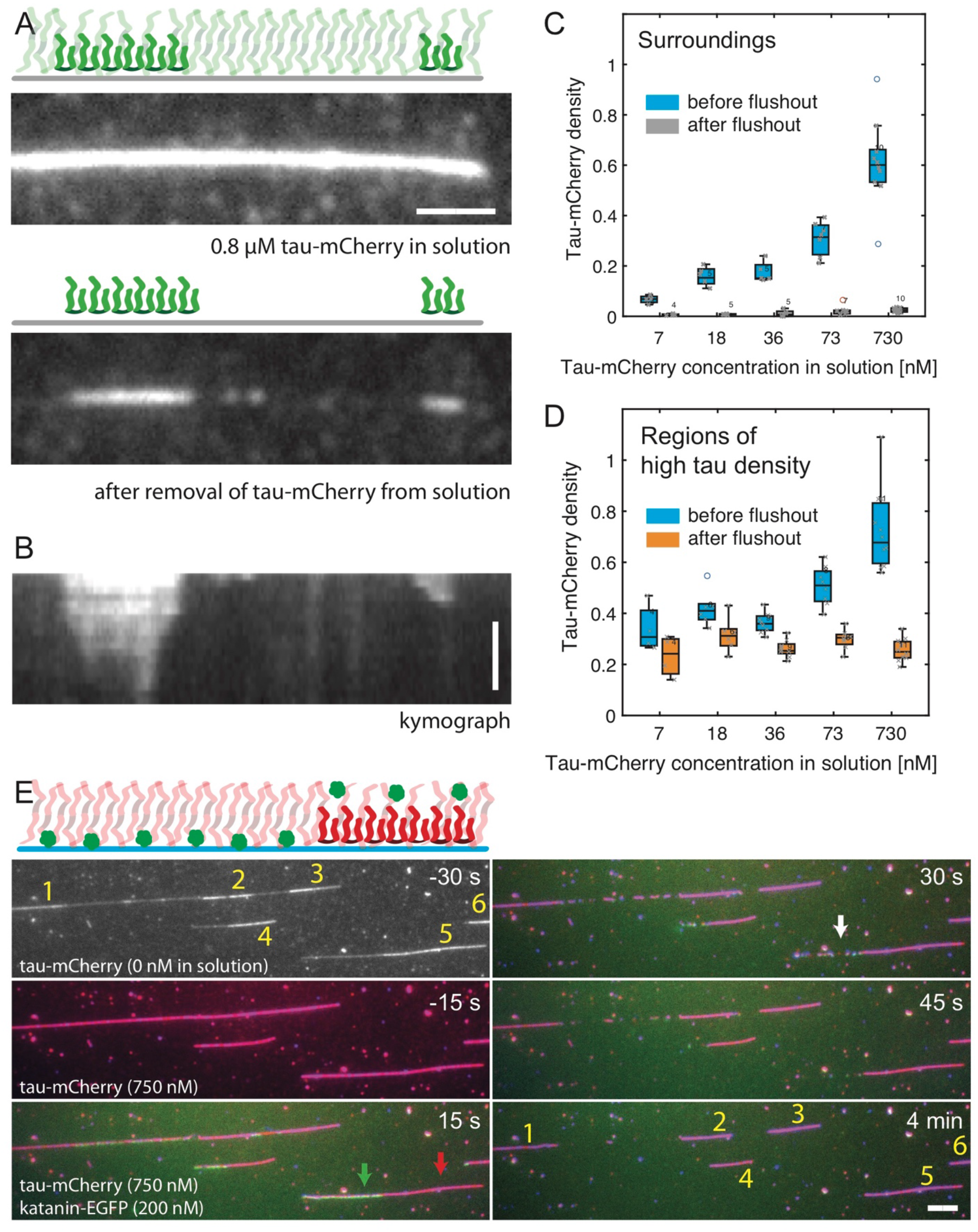
Tau phase which rapidly turns over can colocalize with the islands. **(A)** Fluorescence micrographs showing the initial uniform coverage of the microtubules by tau-mCherry at high (0.8 µM) concentration and the islands as they became visible 30 s after the removal of tau-mCherry from solution. **(B)** Kymograph of the experiment presented in (A) showing the disassembly of the islands. **(C)** Box-plot showing the average densities outside the island regions at various tau-mEGFP concentration prior and after the removal of tau-mEGFP from solution. **(D)** Box-plot showing the average densities in the island regions established at various tau-mEGFP concentration prior and after the removal of tau-mEGFP from solution (data from the same experiment as presented in (C)). **(E)** Fluorescence micrographs showing katanin-GFP-driven (green) severing of Atto-647-microtubules (blue) decorated with tau-mCherry islands (red) formed at 0.8 µM tau-mCherry concentration. The island positions (indicated by numbers) were determined by a brief removal of tau-mCherry from solution (Methods). Katanin is recruited to regions outside of the islands (green arrow) and excluded form the islands (red arrow). Microtubule severing and disassembly occurred initially only in the regions outside of the islands (example indicated by white arrow). Compare to the Fig. 3B. All scale bars, vertical 5 min, horizontal 2 µm.

## Discussion

Taken together, our results show that microtubule-interacting tau can co-exist in two phases: tau molecules that i) explore the microtubule surface by diffusion, or ii) bind stationarily to the microtubule surface forming islands in the pool of the diffusible tau. Tau islands grow by the deposition of tau molecules to the island boundaries in a process reminiscent of epitaxial growth during the formation of thin films (*20*). Island growth is reversible, manifested by the observed island disassembly from their boundaries when tau is removed from solution.

The co-existence of two tau phases on the microtubule can locally regulate the accessibility of the microtubule surface by other MAPs and their activity. It has been shown in vivo that tau protects microtubules from severing by katanin (*7*). Our results reveal an additional facet of this interplay. Tau islands indeed prevent katanin-mediated microtubule severing, whereas katanin can bind to and sever microtubules in regions, where tau interacts diffusibly (even at high densities). This potentially provides a regulatory mechanism for determining where katanin can sever microtubules. It has been reported that tau inhibits kinesin activity in vivo and in vitro (*8-13*). We observed that kinesin-1 can walk processively in the regions of diffusible tau binding outside of the islands. However, islands effectively blocked kinesin-1 binding to the microtubule, which might have consequences for axonal transport. Furthermore, we show that the islands can be regulated by the action of other proteins, such as the super-processive kinesin-8, which was able to walk through and thereby disassemble the islands. This finding suggests that the accessibility of the microtubule surface is governed by the interplay between the affinities of tau molecules constituting the island and the MAPs.

The island-constituting tau molecules turn over slowly and are bound to the microtubule at a characteristic density in the range of 0.25 to 0.45 nm^-1^. This density is similar to the value estimated recently by cryo-electron microscopy (~0.43 nm^-1^), suggesting that islands are monolayers of tau as shown in the respective study (*21*). Further increasing the concentration of tau in solution in our experiments resulted in an increase in the density of tau that rapidly turned over on the microtubule surface, obscuring the islands at physiological tau concentrations. A reason why this pool of tau was not captured by cryo-electron microscopy experiments could be the transiency of the interactions, which might be mediated by the C-termini of tubulin as suggested earlier (*4*). Importantly, this phase of rapidly turning over tau molecules was not able to protect microtubules from katanin severing even at saturating densities at micromolar concentrations. By contrast, islands protected the microtubule from katanin at tau concentrations as low as 20 nM, showing that the presence of an ordered phase of tau is essential for the shielding.

What is the mechanism of the island assembly? In the presence of crowding agents, at micromolar concentrations, tau is known to undergo liquid-liquid phase separation in solution which is underpinned by tau-tau interactions (*22*). As tau-tau interaction on the microtubule have been reported earlier (*23*), it seems plausible that interaction with the microtubule (which effectively restricts the 3-dimensional diffusion of tau in solution to 2-dimensional diffusion along the microtubule surface), enhances the possibility of tau-tau interactions, leading to island-assembly at nanomolar tau concentrations. An alternative explanation for island assembly could be mechanical changes in the microtubule lattice induced by island-constituting tau molecules, leading to increased affinity of tau for neighboring tubulin dimers.

The tau islands described in our study are fundamentally distinct from the tau formations termed “tau patches” reported earlier (*9*). In contrast to patches, the here reported islands are in exchange with the pool of diffusible tau and grow from a nucleation point by the addition of tau to their boundaries. We also note that we did, as suggested earlier (*24*), observe increased binding of tau-mEGFP to regions of highly curved microtubules. However, we observed tau binding to these regions being distinct from islands, both in kinetic behavior and tau density (Fig. S4E, S4F, S4G).

In summary, we show that tau on microtubules separates into distinct phases over a broad range of conditions. Complementary work presented in Tan et al. confirms the existence of tau phase-separation on the microtubule surface and shows its significance for the regulation of cytoplasmic dynein and spastin, extending our results. We hypothesize that the islands of stationary tau can tag specific regions on the microtubules, for example as a readout of differential post-translational modifications of tubulin, rendering the regions on the microtubule surface differentially accessible to other MAPs. Sorting of proteins associated with cytoskeletal filaments can be driven by reciprocal exclusion generated e.g. by geometrical constraints (*25*) or localized binding of different intrinsically disordered proteins (*26*). As tau is not the only disordered protein to have a propensity to undergo liquid-liquid phase separation, we hypothesize that other disordered proteins might be also be able to phase-separate on the microtubule surface, which could add another layer of MAP sorting and regulation on microtubules. It is intriguing to speculate that in neurodegenerative diseases, which involve the gradual dissociation of unstructured proteins from neuronal microtubules, diminished island assembly, triggered e.g. by hyperphosphorylation of tau, could be a cause for various downstream (patho-)physiological effects.

## Acknowledgments

We thank Anna Akhmanova and Kai Jiang for the generous gift of the katanin plasmid, Richard McKenney for feedback and sharing of data, Ilia Zhernov for helpful discussions, Verena Henrichs and Lenka Grycova for help with protein preparation, and Yulia Bobrova for technical support.

## Author contributions

A.H.V. and M.B. first discovered the islands and initiated the project, A.H.V., A.A.H., S.D., Z.L. and M.B. conceived the experiments, A.H.V. generated the tau∆N-mEGFP construct, V.S., J.K., A.H.V. and M.B. generated the proteins, performed and analyzed the experiments, V.S., J.K., S.D., Z.L. and M.B. wrote the manuscript. All authors discussed the results and commented on the manuscript.

## Materials and Methods

### Protein purification

mEGFP or mCherry tagged tau and tau∆N, Kinesin-1, Kip3 and katanin were expressed and purified as described previously (*16*, *22*, *27*, *28*).

### In vitro tau-microtubule binding assay

Microtubules and flow cells were prepared as described previously (*29*). Biotinylated, paclitaxel-stabilized, Atto647-labeled microtubules in BRB80T (80 mM Pipes/KOH pH 6.9, 1 mM MgCl_2_, 1 mM EGTA, 10 µM paclitaxel) were immobilized in a flow chamber using biotin antibodies (Sigma B3640, 20 µg ml^−1^ in PBS). Subsequently, the buffer in the flow cell was exchanged for assay buffer (20 mM HEPES pH 7.2, 1 mM EGTA, 75 mM KCl (unless stated otherwise in the main text), 2 mM MgCl_2_, 1 mM ATP (+Mg), 10 mM dithiothreitol, 0.02 mg/ml casein, 10 µM paclitaxel, 20 mM d-glucose, 0.22 mg/ml glucose oxidase and 20 µg/ml catalase). Then, tau in assay buffer was flushed into the flow cell at the final assay concentration stated in the main text. In experiments including multiple subsequent tau additions, the flow cell was rinsed between each tau addition by high ionic strength buffer (200 mM KCl additional to the assay buffer). To remove tau from solution during experiments presented in Fig. 1 and 4, the chamber was perfused with approximately four-fold amount of the chamber volume using assay buffer not containing tau. For high concentrations of tau (>200 nM), higher volumes (up to ten-fold chamber volume) were used to remove tau. In experiments involving kinesin-8, katanin, or kinesin-1, islands were first preformed before the respective protein was added to solution (keeping the tau concentration constant). For the katanin experiment at elevated tau concentration (Fig. 4E) microtubules were first incubated with 0.8 µM tau-mCherry for 5 minutes. Tau-mCherry was then removed from the measurement chamber for a brief period of time (less then 1 minute) to note the position of the islands (which were obscured by the high tau-mCherry density in the island surroundings). Tau-mCherry was then again added at 0.8 µM. After 5 minutes 215 nM katanin-GFP was added to the solution (while keeping the tau concentration 0.8 µM). All experiments were performed at room temperature.

### Imaging

Atto647-labeled microtubules, mCherry- and mEGFP-labeled tau were visualized sequentially by switching between the Cy5, TRITC and GFP channels (Chroma filter-cubes) using Nikon-Ti E microscope equipped with 100x Nikon TIRF objective and either Hamamatsu Orca Flash 4.0 sCMOS or Andor iXon EMCCD cameras. The acquisition rate varied between 1 frame per 30 ms to 1 frame per 30 seconds depending on the particular experiment and is indicated in the corresponding figure. Imaging conditions in experiments used for quantitative estimation of kinetic parameters (Figs. 1E, 1G, S1C, S2A, 4C, 4D, S4B) were set such that photo-bleaching effects were negligible (< 2 % fluorescent intensity loss during the experiment). Experiments were performed over several months by three independent experimentalists. No data was excluded from the study.

### Image analysis

Data was analyzed using FIJI (*30*) and custom written Matlab (Mathworks) routines. Kymographs (KymographBuilder plugin, custom-modified to compute integrated intensity instead of finding the maximum intensity) along the microtubule length were used to read out the eMGFP or mCherry fluorescent signal and to estimate the integrated signal intensity of mEGFP- or mCherry-labeled tau bound to the microtubule (if necessary, time series were drift-corrected with FIESTA (*31*)). The mEGFP signal in regions directly adjacent to the microtubule was estimated in the same way, smoothed with a moving median along the microtubule length and subtracted as background signal. The kymograph pixels were then manually categorized according to the type of microtubule region they covered (island, curved microtubule, regions surrounding the islands). Integrated intensity traces for each region type were then computed for each frame by taking the mean of the corresponding kymograph pixels. The density of mEGFP- or mCherry-labeled tau bound to the microtubule was then estimated by dividing the integrated intensity by estimated intensity per single molecule times unit length. The corresponding traces underlie all figures where Tau densities are given, with N being the number of traces. Single numbers for any given condition were obtained by taking the median of the relevant frames of each density trace and averaging the obtained median values.

Curve fitting was performed with the Matlab function *fit*, whereas each density trace was fitted separately. The frames taken before flushing in different solutions had been averaged, and the resulting value had been assigned a weight double the size of the other points in the trace.

Fluorescent signal of a single fluorescent molecule (mEGFP or mCherry) was determined by generating intensity time-traces of single mEGFP- or mCherry-labeled kinesin-1 molecules tightly bound to the microtubule in presence of AMP-PNP (in the absence of ATP) and estimating the height of the occurring bleaching steps. The number of steps was first estimated by eye, and this number was used as input for the *findchangepoints* function of Matlab to determine the position of the steps (by detection of significant changes of the mean value). To yield the intensity per single molecule, the heights of these steps were averaged (after removing outliers via Hampel filtering). The number of averaged steps was at least 10 per estimate. Photo-bleaching rates (at the given imaging conditions, on the day of the experiment) were determined using the experimental setup described above. Instead of single molecule intensities, here the integrated intensity of all labeled kinesin-1 molecules was measured.

---

**Fig. S1.**
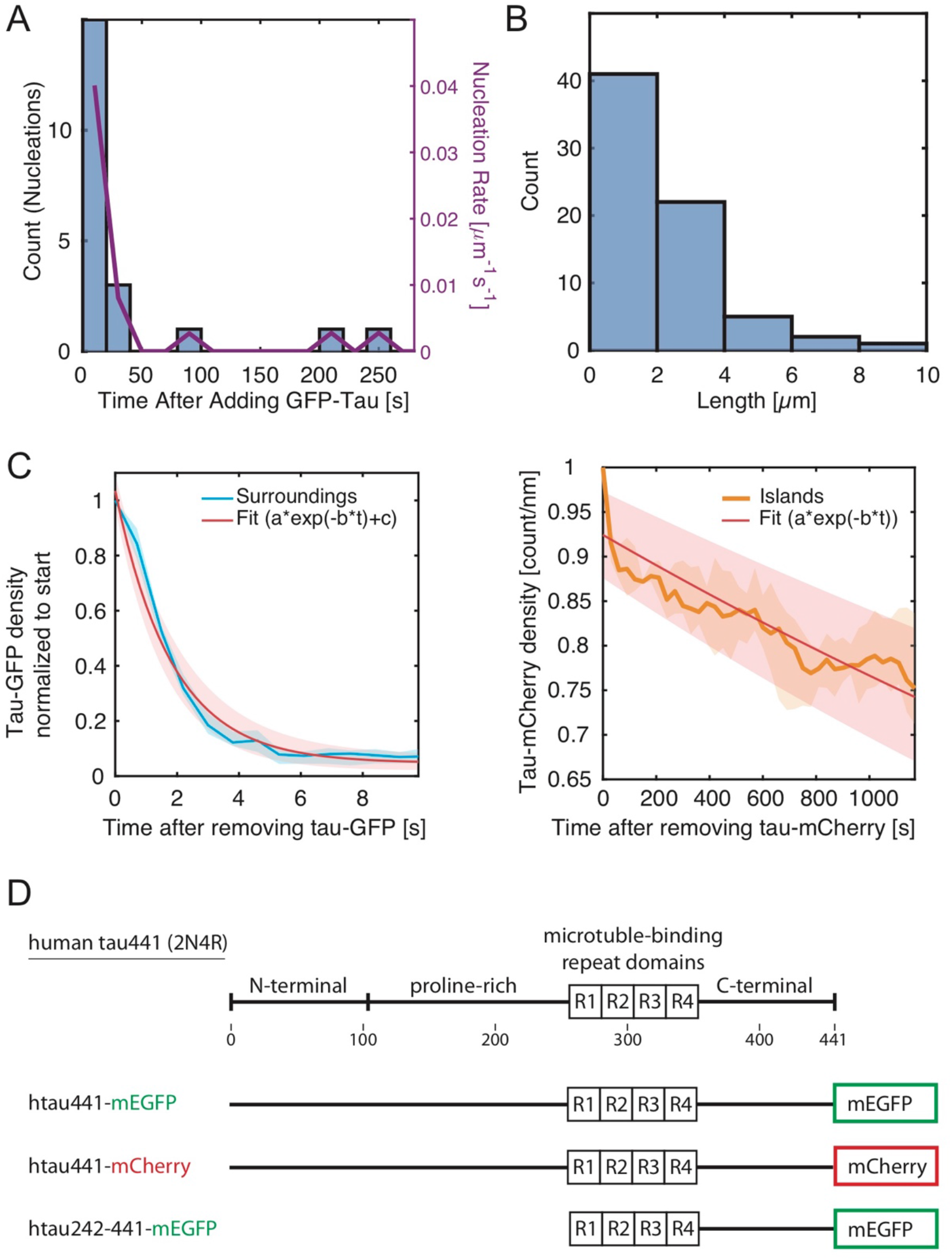
**(A)** Distribution of time between the addition of 20 nM tau-mEGFP and the nucleation of the islands. **(B)** Distribution of the island lengths 15 minutes after the addition of tau-mEGFP. **(C)** The tau-mEGFP density within and outside the islands after the removal of tau-mEGFP from solution as shown in the Fig.1G. Single exponential fits to the data (red lines) yield the tau-mEGFP dwell time outside of the island of 1.9 ± 0.2 s and inside of the island of 93 ± 20 min. **(D)** Schematics showing the constructs used in this study.

**Fig. S2.**
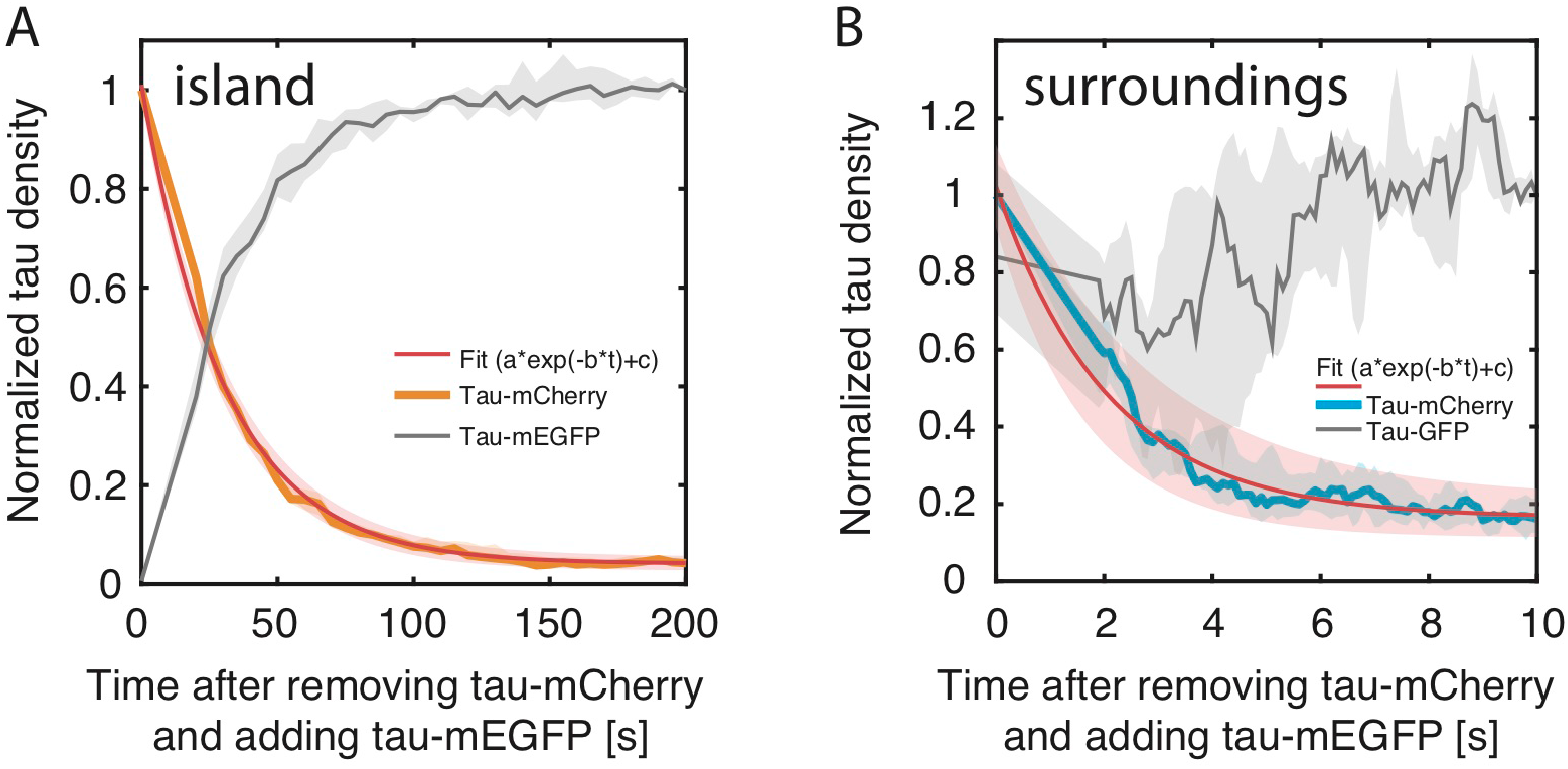
**(A)** Plot of normalized tau-mEGFP and tau-mCherry density within and outside of the island regions after the tau-mCherry - tau-mEGFP exchange. Single exponential fits to the data (red lines) yield the tau-mCherry dwell time outside of the island of 2.2 ± 0.5 and inside of the island of 31 ± 2. The photo-bleaching during this experiment was negligible (Methods).

**Fig. S3.**
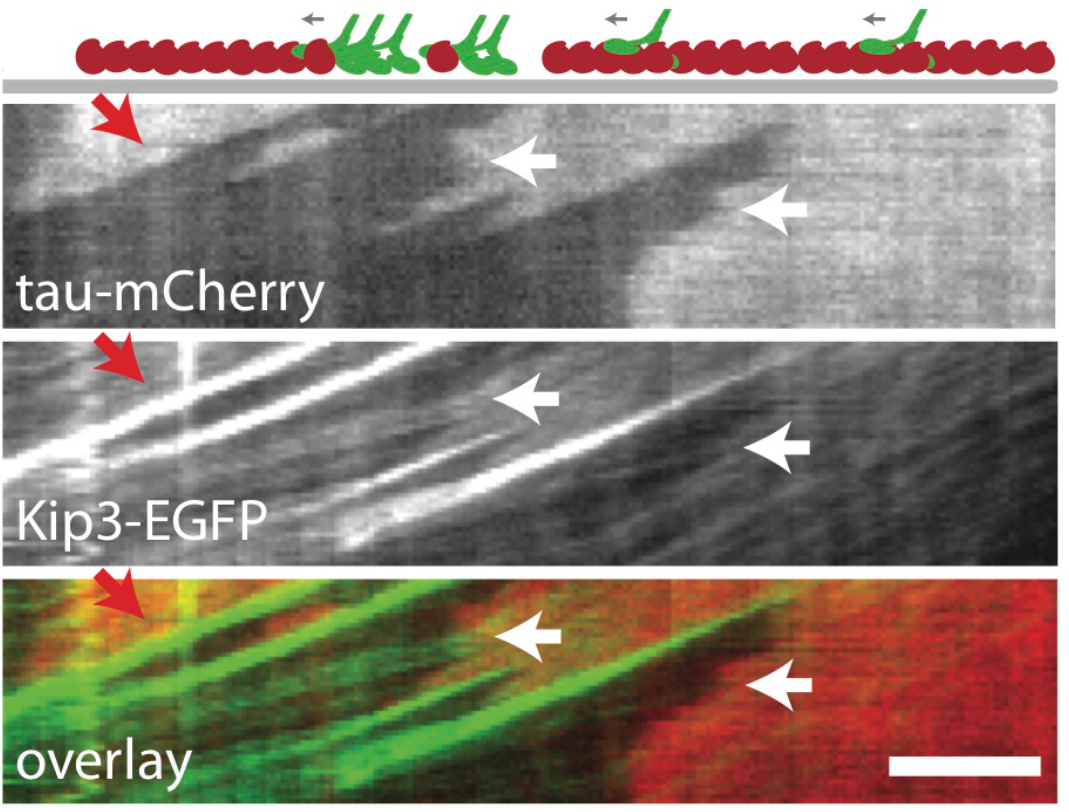
**(A)** Multichannel fluorescence kymograph showing the motion of kip3-EGFP (green) within and outside the island regions (red). Red arrow indicates the accumulation of kip3-EGFP in front of the island. White arrows indicate kip3-EGFP molecules speeding up as they leave the island. Scale bar 2 µm.

**Fig. S4.**
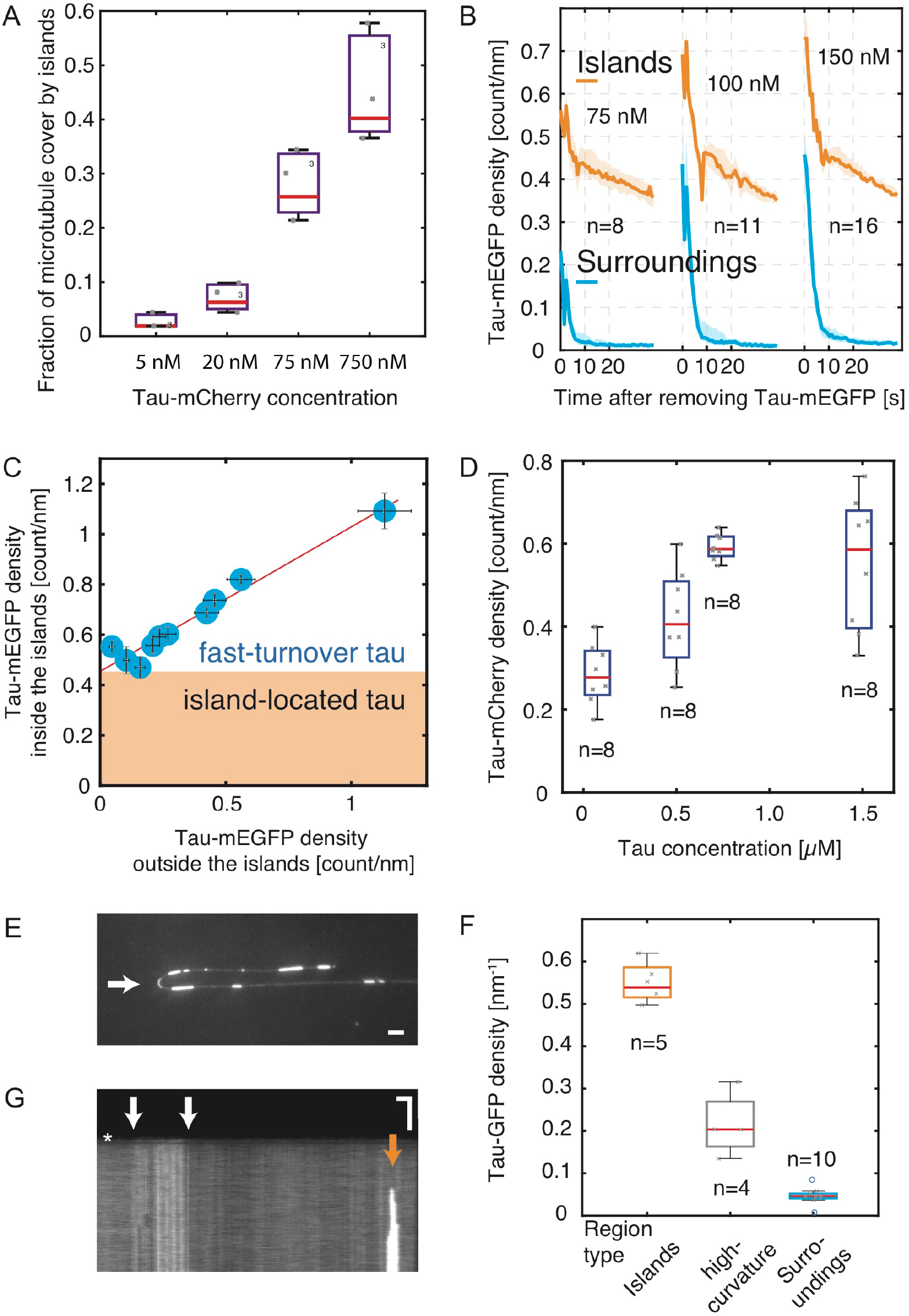
**(A)** Fraction of the microtubule length covered by islands 5 minutes after the addition of tau-mEGFP as a function of the tau-mCherry concentration in solution. **(B)** Time-trace of tau-mEGFP density outside and within the island regions during the subsequent cycles of i) additon of increasing concentrations of tau-mEGFP followed by ii) the removal of tau-mEGFP from solution. Experiment such as presented in Fig. 4C,D. **(C)** At tau-mEGFP concentration range of 20 nM to 0.8 µM, tau-mEGFP density within the islands scales linearly with the tau-mEGFP density outside of the islands and is offset by the factor of 0.45, which is the density of the slow-turnover tau-mEGFP population observed in Fig. 4D. Data from experiment presented in the Fig. 4C,D. **(D)** Density of tau-mEGFP within and outside of the island regions saturates at high (µM) tau-mEGFP concentrations. **(E)** Fluorescence micrograph showing tau-mEGFP signal on microtubules. Tau islands have higher tau-mEGFP density than the tau-mEGFP regions localizing to microtubule curves (white arrow). See also quantification in (F). **(F)** Boxplot of densities of tau-mEGFP outside of the islands, in the islands and in the regions of high curvature showing that the density in the curves is significantly lower than in the islands localized along straight microtubules. **(G)** Kymograph showing the uniform increase of tau-mEGFP density along the whole region of the microtubule curve (white arrows) upon the addition of tau-mEGFP in solution. This is in strong contrast to islands localized on straight microtubules, which grew at their boundaries (orange arrow). Combined with data presented in (E and F) these results suggest that the tau formations on highly curved microtubules are different from the islands on straight microtubules discussed in Fig. 1. Scale bars, vertical 10 min, horizontal 2 µm.

**Movie S1.** Formation of islands of high tau-mEGFP density on Atto647-labeled microtubule (not shown) after the addition of 20 nM tau-mEGFP. Experiment presented in Fig. 1C.

**Movie S2.** Disassembly of the tau-mEGFP islands upon the removal of tau-mEGFP from solution. Experiment presented in Fig. 1F (the frames before removing tau from solution are not shown in the kymograph).

**Movie S3.** Tau turnover within and outside of the patches visualized after the exchange of 20 nM tau-mCherry (red) for 20 nM tau-mEGFP (green). Experiment presented in Fig. 2A.

**Movie S4**. Katanin-GFP-mediated microtubule severing and subsequent microtubule disassembly occurred initially only in the regions outside of the islands. On the longer time scales also the island-covered parts of the microtubules started disassembling from their boundaries. Experiment presented in Fig. 3B.

**Movie S5.** Kip3-EGFP (green) moves both, in the low tau-mEGFP density regions (dim red) and in the islands (bright red). Kip3-GFP that accumulated in front of a tau-mCherry island caused enhanced unbinding of tau-mCherry from the boundary of the island leading to the complete removal of the island. Experiment presented in Fig. 3C, D.

**Movie S6.** Tau-mCherry at a concentration of 0.8 µM covers microtubules uniformly. With addition of Katanin-GFP, we, as in Movie S4, observed Katanin-GFP binding and microtubule disassembly predominantly in regions which are not shielded by tau islands. Experiment presented in Fig. 4E.

